# High-Accuracy RNA Integrity Definition for Unbiased Transcriptome Comparisons with INDEGRA

**DOI:** 10.1101/2024.12.12.627949

**Authors:** Alice Cleynen, Agin Ravindran, Aditya Sethi, Bhavika Kumar, Tanya Javaid, Shafi Mahmud, Katrina Woodward, Helaine Graziele Santos Vieira, Minna-Liisa Änkö, Robert Weatheritt, Eduardo Eyras, Stéphane Robin, Nikolay Shirokikh

**Affiliations:** The Shine-Dalgarno Centre for RNA Innovation, The John Curtin School of Medical Research, ANU, Canberra, Australia; CNRS, Université de Montpellier, Montpellier, France; Garvan Institute of Medical Research, Darlinghurst, Australia; Tampere University, Tampere, Finland and Hudson Institute of Medical Research, Clayton, Australia; Sorbonne Université, Paris, France

## Abstract

RNA sample integrity variability introduces biases and obscures natural RNA degradation, posing a significant challenge in transcriptomics. To address this, we developed the Direct Transcriptome Integrity (DTI) measure, a universal and robust RNA integrity metric based on nanopore sequencing. By accurately modeling RNA fragmentation, DTI provides a reliable assessment of sample quality. Integrated into the INDEGRA package (freely available at https://github.com/Arnaroo/INDEGRA), we provide tools to correct false discoveries and enable precise differential expression and RNA degradation analyses, even for challenging sample types.

INDEGRA software can be used to accurately measure RNA DTI stability metric, isolate biological component of RNA degradation from technical biases, compare biological RNA stability transcriptome-wide and suppress false degradation-induced differential gene expression hits to allow broad comparisons across samples of different quality

DTI offers a straightforward and accurate method for assessing RNA degradation, characterizing both overall sample integrity and transcript-specific degradation rates using direct RNA sequencing (DRS) data. Calculated through INDEGRA, DTI reveals inter- and intra-transcript variability in degradation, while INDEGRA separates RNA degradation from mapping inaccuracies, and connects degradation profiles to RNA fragmentation rates. By leveraging INDEGRA, researchers can minimize false differential transcript abundance findings caused by variations in overall sample integrity, while preserving genuine transcript-specific differences in stability and degradation.

INDEGRA supports integration with widely used differential transcript abundance tools like DESeq2, limma-voom, and edgeR, enabling seamless analysis pipelines. INDEGRA enhances the accuracy and reliability of RNA quantification in high-throughput data and simplifies comparisons across diverse transcriptomic datasets, including those derived from different tissues, species, or experimental protocols.

## Introduction

RNA carries immense amounts of information about gene expression and the exact cell’s state but is a subject to a wide range of decay processes^1–5^. Every cellular transcript has a distinct half-life, which extends for messenger (m)RNA from about 10-15 minutes (*e*.*g*. in many proto-oncogene mRNAs) to several hours (globin mRNAs)^6^ in a typical mammalian cell. mRNA turnover naturally replenishes damaged, translationally stuck or unwanted RNA with the new types, and aids in reprogramming along with the internal (*e*.*g*. cell cycle, signaling) or external (*e*.*g*. nutrients, stress) stimuli^4,7,8^. Of the notable intracellular RNA decay mechanisms, decapping by the enzymes of the DCP family followed by the 3′→5′ deadenylation by CCR4-Not complex and 5′→3′ degradation driven by XRN-family proteins, together with the 3′→5′ degradation by the exosome complex are the major routes of cytosolic (and sometimes nuclear) mRNA degradation (**Figure 1**)^9^. While the exosome can degrade various types of RNA including ribosomal RNA, the 5′→3′ exoribonucleolytic mRNA decay is considered to be overall a dominant cytosolic decay mechanism of the eukaryotic cells^5,10^. More specialized pathways include nonsense-mediated decay (mediated by UPF family of proteins) or microRNA-induced degradation (mediated by Argonaute family of proteins), which act on predefined sets of targets^2,5,9,11^.

**Figure 1.**
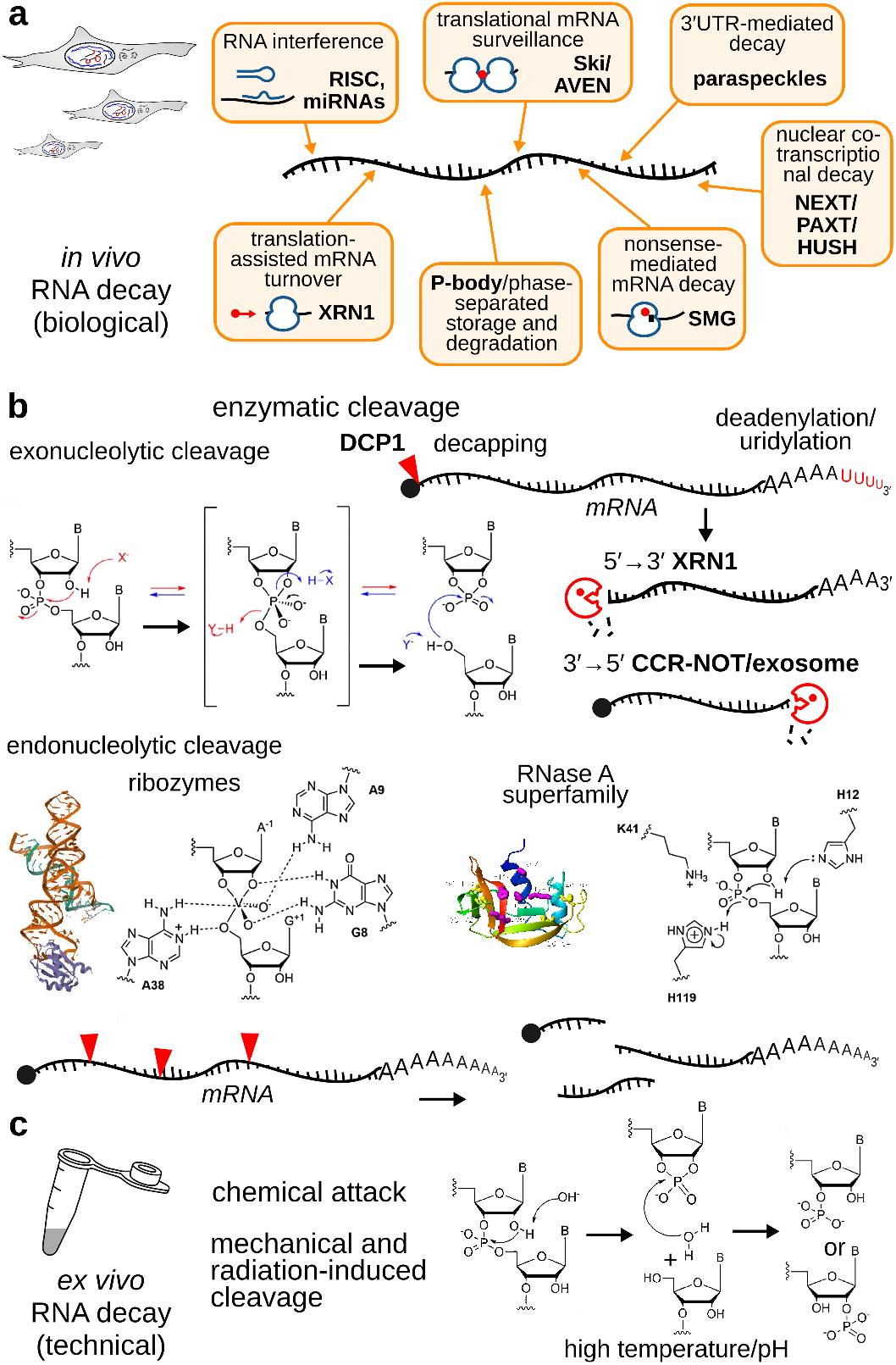
Pathways and mechanisms of RNA degradation. **(a)** Schematic depicting various pathways of degradation *in vivo* (‘biological’ decay). **(b)** Mechanisms of enzymatic cleavage. **(c)** Less specific and commonly *ex vivo*-encountered mechanisms (‘technical’ decay).

RNA can also be degraded extracellularly^8^ or as a result of adversary non-natural effects on the cells, including during sample handling. This technical degradation is often driven by the extremely stable and ubiquitous endonucleolytic enzymes of RNase A superfamily and similar, as well as chemically-induced hydrolysis at higher pH, temperatures and in the presence of multivalent and especially transition metals with different oxidation states^12,13^.

RNA sample integrity is critically important to the accurate measurement of differential transcript abundance measurement and other RNA feature quantification^14^. Yet it is not always possible to have samples with matching integrity levels, and the degradation assessment may be insufficiently convenient or biased towards certain RNA types.

Several useful tools have been developed to estimate RNA integrity, of which RNA Integrity Index (RIN)^15^, DV200^16^, Transcript Integrity Index (TIN and TII)^17,18^ and mRIN^19^ are often employed. However, RIN is closed-source and uses machine learning assessment of electrophoretically separated samples mostly accounting for the major total RNA constituents, largely ribosomal RNA, whereas DV200 also interprets electrophoresis outcomes and is based on the rate of fragments longer than 200 nt. These tools are also not appropriate for many sample types, such as those lacking ribosomal RNA or having large fraction of shorter RNAs, and do not provide transcript-level information. TIN, TII and mRIN use short-read sequencing mapping and read distribution tests to assess degradation. However, these tools employ amplified signal with high coverage biases, cannot link to the physical degradation rate and do not account well for the RNA isoforms.

We thus set out to develop a solution that would address the isoform-resolved quantification and amplification bias problems, while being robust and simple to use. We reasoned that to achieve these objectives, we first need to accurately measure the RNA degradation rate, and then correct differently-degraded samples in a uniform and unbiased way, to enable identification of truly differentially degraded transcripts.

## Results

### Choice of Direct RNA Sequencing to study degradation

To accurately measure the degradation, we have turned to direct RNA sequencing (DRS) data that provide a unique opportunity to investigate RNA molecules in their *in vivo* configuration, in full length, allowing isoform attribution, with minimal library construction bias and without amplification artifacts^20,21^ (**Figure 2a**). Due to the accessibility of the sequencing costs, information about the native primary structure of RNA and ease of sequencing, DRS experiments have gained high popularity.

**Figure 2.**
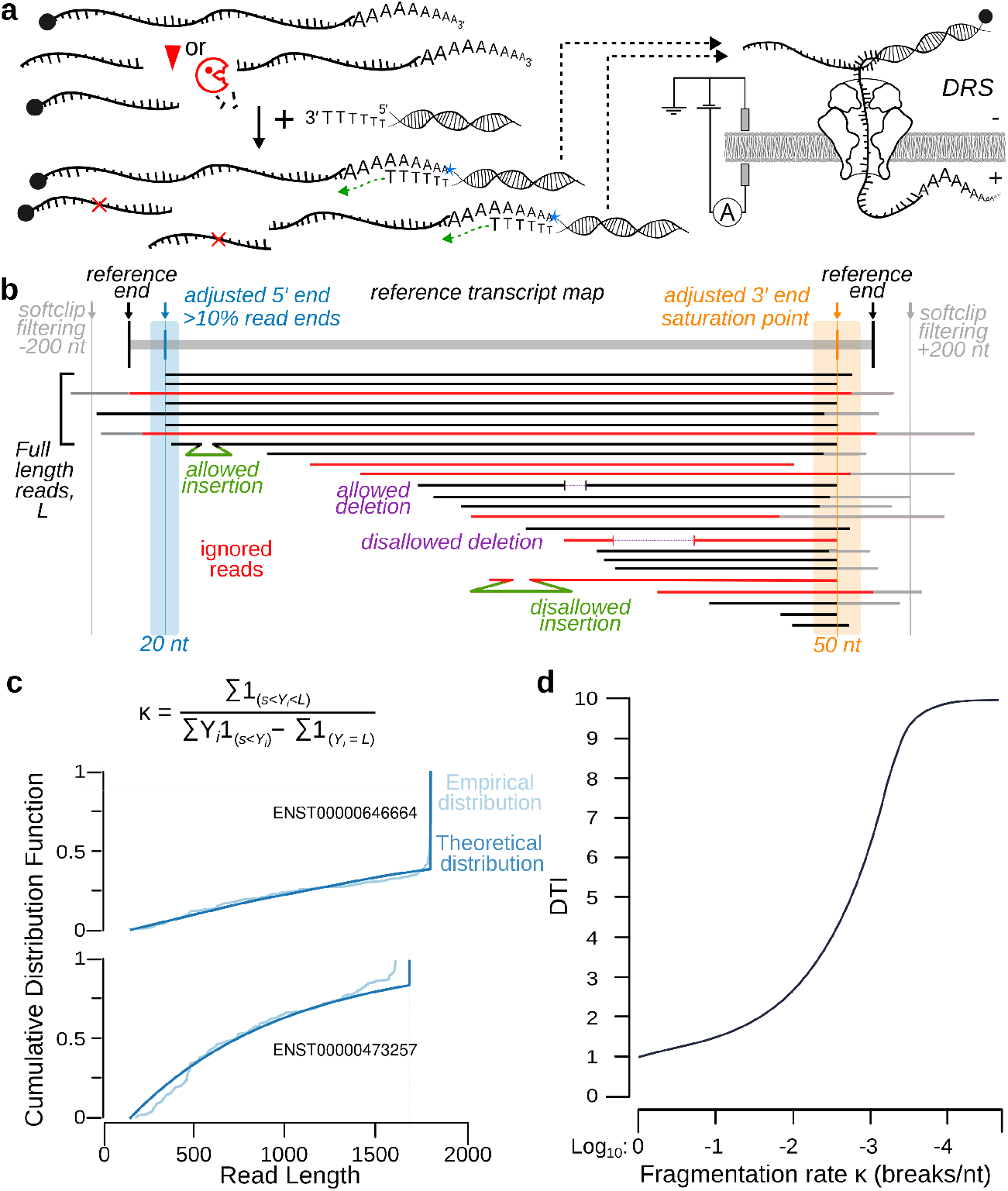
The INDEGRA software for evaluation of differentially-degraded RNA samples and transcripts, and the universal and robust Direct Transcript Integrity (DTI) index. **(a)** Capture of the native, unamplified 3′-polyadenylated RNA and its decay fragments by the default Oxford Nanopore technologies DRS protocol. **(b)** Schematic illustrating the main post-alignment read filtering and transcript end definition steps employed by INDEGRA. **(c)** Calculation of the total fragmentation rate κ (cuts/nt) is shown, together with the examples of model fit for two ACTB isoforms from the human cerebellum sample. **(d)** Conversion of the fragmentation rate κ into RIN-like, per-transcript or per-sample Direct Transcript Integrity (DTI) index.

To discard a majority of bias that would originate from approximate annotation or alignment errors, we introduce an extra layer of filtering by discarding all reads with large insertions, deletions or softclip lengths, and propose a re-annotation of the 3’ and 5’ ends of the transcript from which additional filtering is performed (**Figure 2b** and **Materials and Methods**).

### Mathematical Modeling of degradation

We argued that random fragmentation is a least assumptive representation of RNA degradation, and can be universally applied to most cases of biological or technical RNA decay. We modeled random fragmentation as a homogeneous Bernoulli process^22^ of rate κ, where κ is interpreted as the probability that a fragmentation event occurs between any two nucleotides.

This implies that in naturally polyadenylated samples, the theoretical distribution of read lengths is a truncated geometric distribution. Thus, for any transcript of size L, the probability of observing a read of length l is (1 – κ)^{L – 1}^ for l = L, and κ × (1-κ)^{l – 1}^ for l < L. In particular, for any l smaller than L, this probability does not depend on the transcript length.

The maximum likelihood estimator of κ is a simple relationship between number of full-length reads, non full-length reads, and the sum of read length (**Figure 2c**). We account for characterized DRS bias towards shorter molecules by computing read-length distributions conditionally on observing only reads longer than a given threshold s (user-tunable, default s = 150 nt was set based on the experience with DRS). The rate of fragmentation can then be measured by a maximum likelihood estimator of 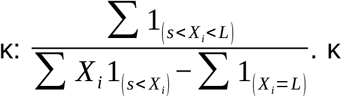 is the ratio of the number of non full-length reads to the total read length less number of full-length read (retaining only reads with length greater than s).

### Introduction of the DTI metric

We then transformed κ to a new per-transcript metric termed Direct Transcript Integrity (^*t*^DTI), which we designed with a behavior similar to RIN whereby ^*t*^DTI of 1 is the maximum degradation rate (cut at every nucleotide), ^*t*^DTI of 10 has the minimal degradation rate (no fragmentation at all), while the linear part of the DTI response falls onto the meaningful degradation rates of one cut within every 500 to 2,000 nucleotides (**Figure 2d**). This is done via the mapping function

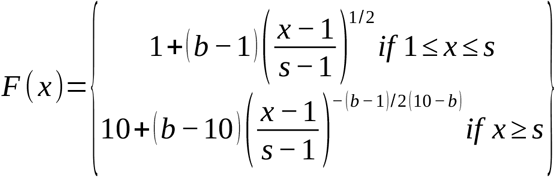

Then, transcript-wise ^*t*^DTI is defined as F(κ), and per-sample DTI is computed as the median of all ^*t*^DTI.

### Testing for deviation to random degradation

A per-transcript Chi-square goodness of fit test is used to compare the empirical distribution of read-lengths to the theoretical distribution under random fragmentation with rate κ. To make the test more robust, k bins of size l = 30 were considered, and the statistic was computed as follows: 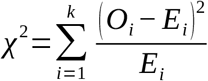, where O_j_ and E_j_ were respectively the observed and expected numbers of reads with lengths within window j. Under the random fragmentation hypothesis, Chi-square follows a standard chi-square distribution with k – 1 degrees of freedom. Adjustment for multiple testing is then performed with Benjamini-Hochberg^23^ procedure.

### Validation of the DTI model

To validate and calibrate DTI, we used controlled magnesium-induced random fragmentation of a set of degradation reference transcript spike-ins of different length and unique sequence (i1, i3, i4) and HEK293 total RNA, and rigorously compared their DTI-assessed integrity with fragmentation determined by electrophoretic separation (**Figure 3a**). The fragmentation rate and negligible effects on degradation during sequencing were controlled with i1, i3 included before and i4 after the fragmentation. This allowed us to confirm direct relation of ^*t*^DTI to the physical degradation rate. (**Figure 3b**).

**Figure 3.**
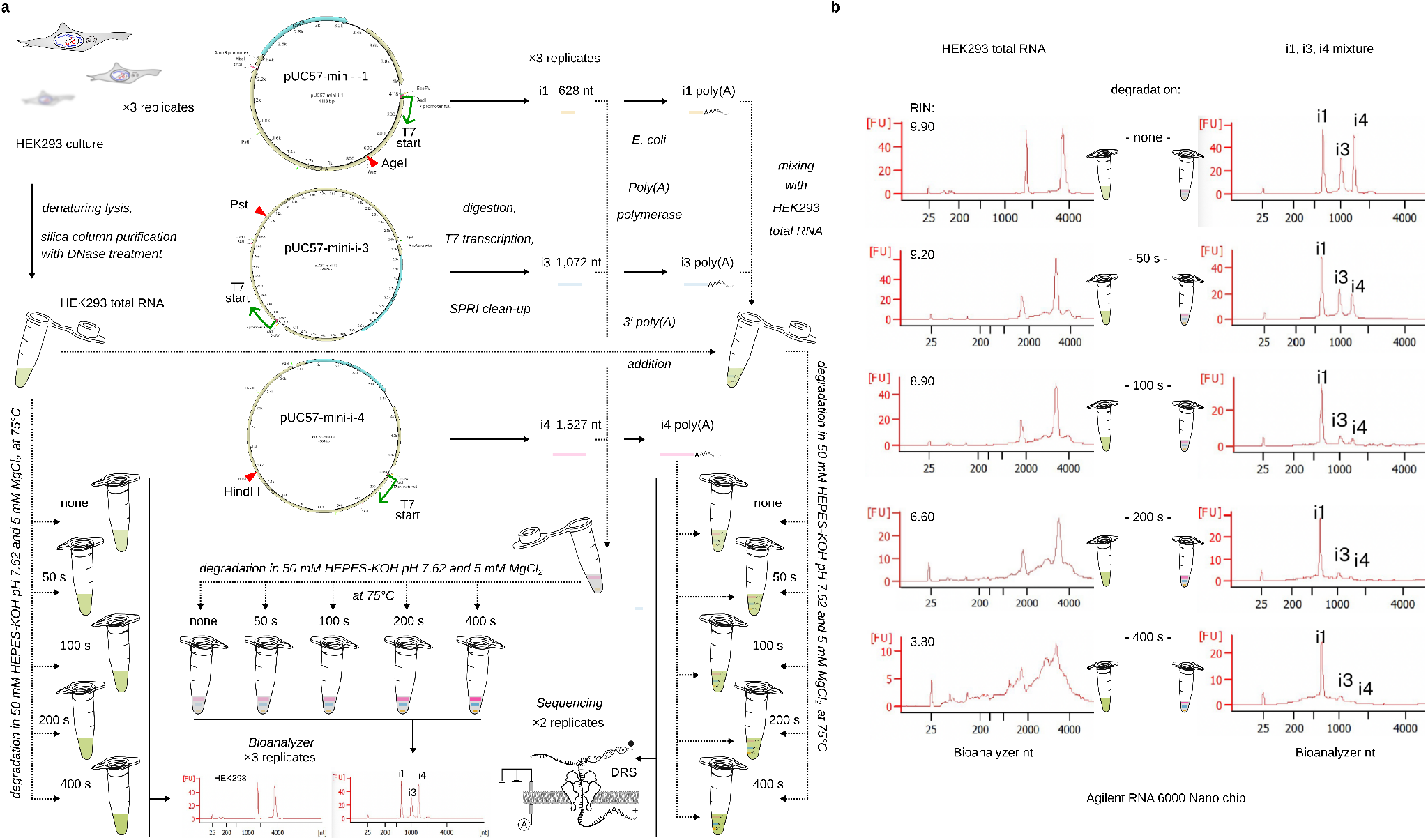
Controlled RNA degradation and DTI validation approach and data. **(a)** Schematic of experimental steps used to set up controlled magnesium ion-based degradation in native RNA and i-type synthetic RNAs of a different length. Illustrated are collection of the total HEK293T total RNA (left) and run-off transcription-based generation of i-type synthetic RNAs (right) from pUC57-mini-i plasmids. Total RNA extraction was conducted using PureLink RNA Mini Kit based on silica columns with DNase treatment (Thermo Fisher Scientific) and used as is, whereas i-type RNAs were cleaned-up using SPRI isolation and used in the form of a mixture, or additionally 3′-polyadenylated with Escherichia coli poly(A) polymerase before use. For the biophysical (electrophiretic) assessment of the fragmentation extent, the combined non-polyadenylated i-type synthetic RNAs and HEK293T total RNA were fragmented in separate experiments using temperature-induced degradation in 5 mM magnesium chloride solution for 0, 50, 100, 200 and 400 seconds. At each condition, the degradation profiles were analyzed qualitatively and quantitatively using Bioanalyzer 2100 runs with RNA 6000 Nano chips (Agilent). For the DRS experiments, HEK293 total RNA was spiked-in by the i-type RNA mixture of pre-polyadenylated i1 and i3 RNAs, followed by temperature-induced degradation in 5 mM magnesium chloride solution for 0, 50, 100, 200 and 400 seconds. The resultant material was then combined with the undegraded pre-polyadenylated i4, and subjected to direct RNA nanopore sequencing (DRS). Two biological replicates were used in all cases. **(b)** Bioanalyzer 2100 (Agilent) RNA 6000 Nano fluorescence traces of i-type RNA mixtures (i1, i3 and i4; left column) and total HEK293 RNA (right column) subjected to different extent of magnesium ion-induced degradation. The traces include results from intact control samples (top, no high-temperature incubation) and samples with the increasing degradation extent (towards the bottom; 75°C incubation for the indicated time). For each degradation time point, a total of three biological replicates were run and the reported RIN values for HEK293 material is an average of these replicates.

Furthermore, DTI showed excellent correlation with RIN for 63 total RNA samples from various species and tissues with well-represented rRNA we have tested, further validating the metric (**Figure 4a**). Importantly, while DTI median of the entire transcriptome correlates with the RIN without weighting in transcript abundances, per-transcript ^*t*^DTI shows tissue-specific distribution, high variability (**Figure 4b**) and reveals significant degradation differences between RNA isoforms (**Figure 4c**).

**Figure 4.**
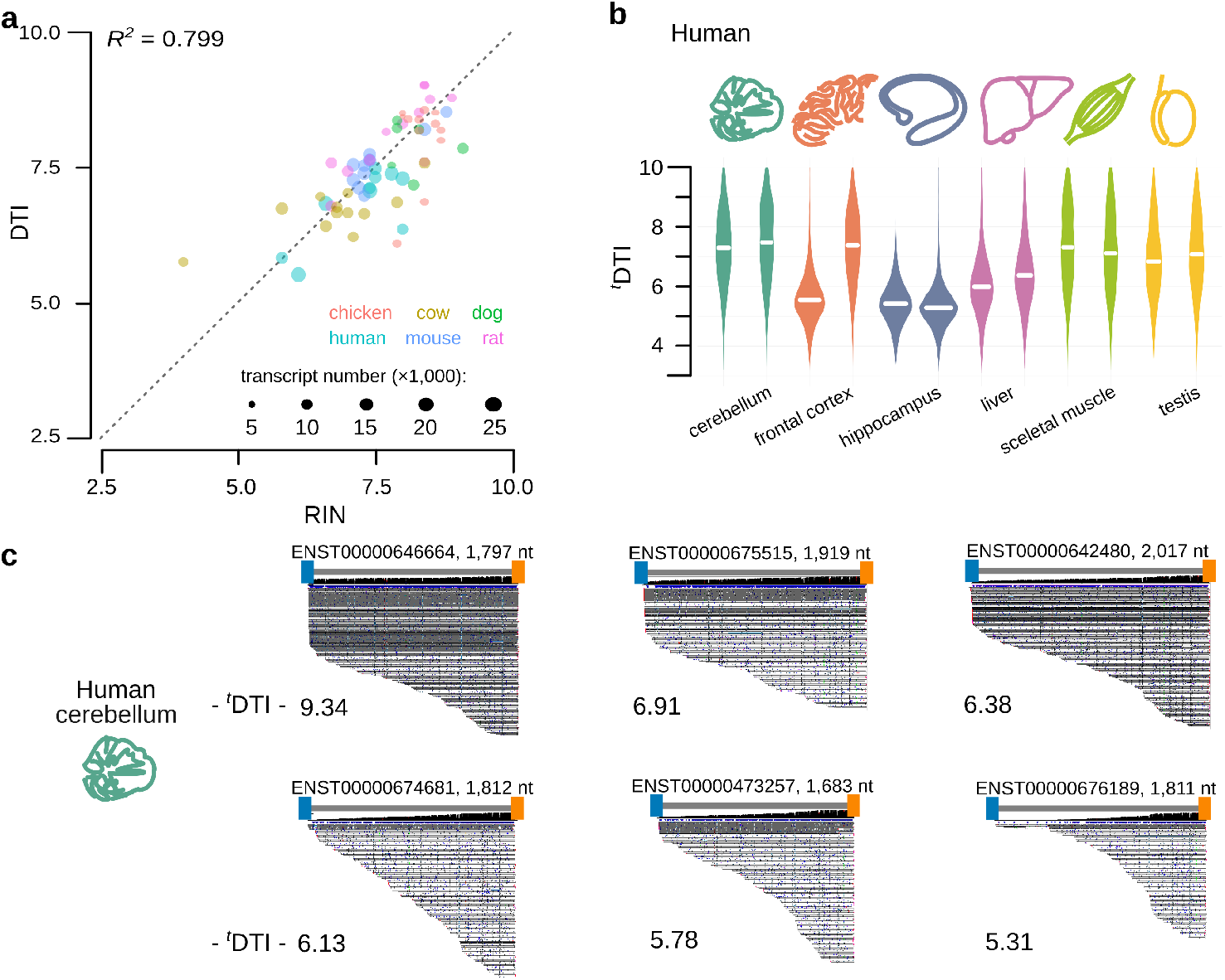
Application of INDEGRA to assess RNA sample and transcript-specific integrity. **(a)** Observed relationship between the sample DTI and RIN across a replicated DRS dataset derived from 6 species and 6 tissues of vertebrates. **(c)** Highly tissue-specific transcript-wise ^*t*^DTI distribution pointing at biologically-important mRNA stability differences across human tissues. **(d)** Discovery of isoform-specific stability using select examples of β-actin (*ACTB*) gene transcripts detected in human cerebellum samples.

### Deconvolution of biological and technical degradation

To exploit the DTI power in dissecting biological transcript-selective degradation rate, we employed Bayesian modeling and deconvolved biological and technical degradation based on the the read length distribution. Here we reasoned that any technical degradation should occur after biological degradation, allowing to build a model segregating these and highlight transcripts with unusual stability or instability *in vivo*, even across samples which have overall different integrity (**Figure 5a**). We thus constructed a _δ_^*t*^DTI test for differences in biological degradation at the transcript level, which can detect condition-specific RNA decay even in samples of uneven quality (**Figure 5b**). The test is based on the posterior probability of same biological degradation rate occurring in two independent samples, see **Materials and Methods**. Removing the contribution of technical degradation with _δ_^*t*^DTI, we can identify groups of universally stable or unstable transcripts, and those stable or unstable in select tissues, revealing species- and tissue-specific re-tuning of the transcriptome turnover (**Figure 5c**).

**Figure 5.**
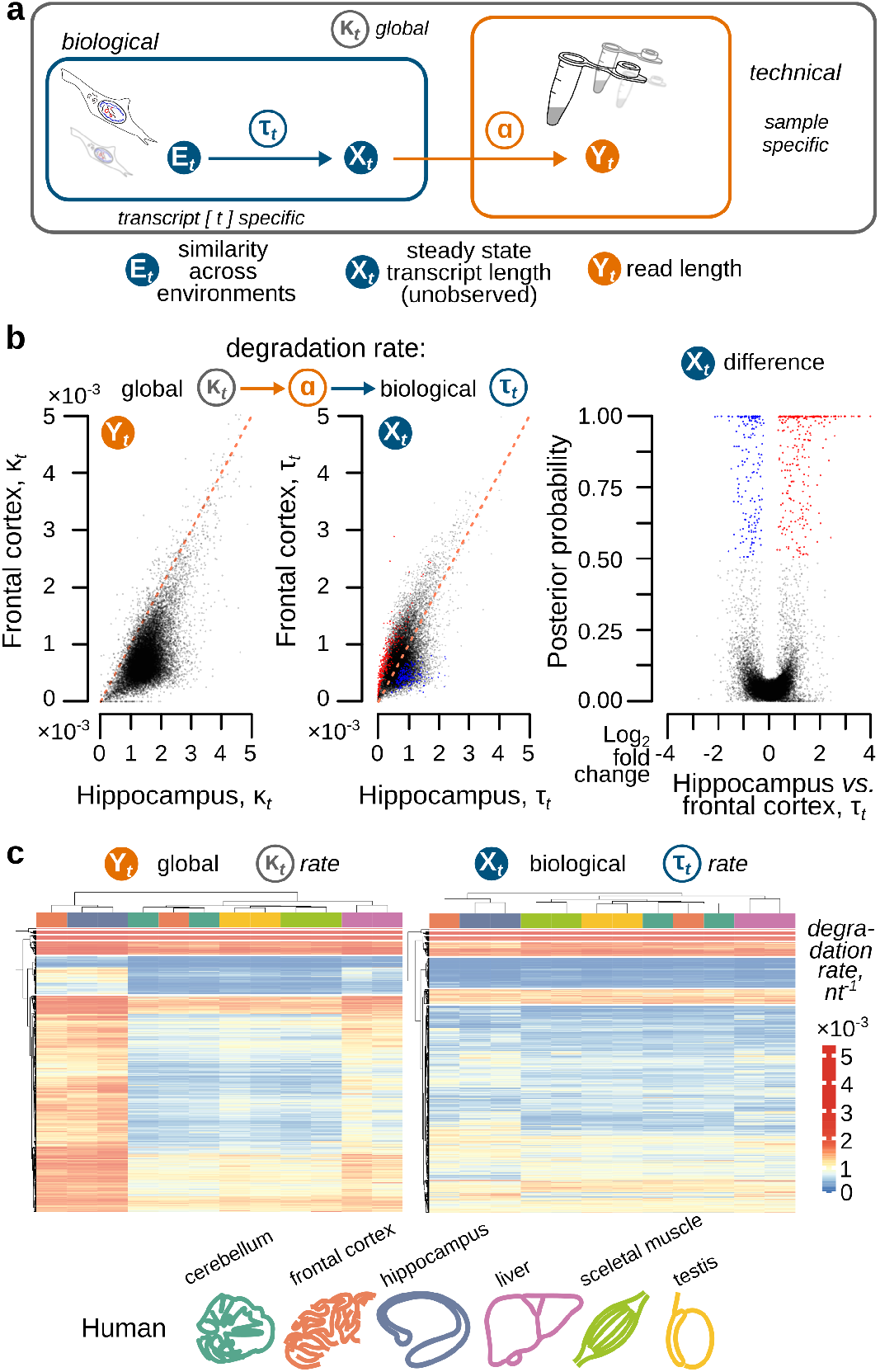
Application of INDEGRA to detect and neutralize technical degradation component, and reveal biological stability differences between samples and transcripts. **(a)** Schematic of the Bayesian approach to deconvolve technical (α) and biological (τ _*t*_) degradation across RNA samples employed in INDEGRA. **(b)** Correcting for technical degradation to reveal biological transcript stability differences across human hippocampal and frontal cortex total RNA samples. Shown are total (κ_*t*_) and biological (τ_*t*_) degradation (as cuts per nucleotide). **(c)** Rectifying biological stability differences to broadly investigate degradation preferences across human tissues with INDEGRA. κ_*t*_ values (left heatmap) are compared with the inferred biological degradation τ_*t*_ values (right plot).

To rigorously validate _δ_^*t*^DTI, we used accurately calibrated low (0.995 _δ_DTI, 0.500 _δ_RIN) mild (1.275 _δ_DTI, 0.850 _δ_RIN) and strong (2.085 _δ_DTI, 6.000 _δ_RIN) randomly-fragmented versions of the total HEK293 transcriptome with the original DTI of 6.855 (RIN of 9.900). Here we show comparing the undegraded samples to each of the degraded conditions that the number of transcripts with significant difference in overall degradation rate drastically increases as the magnesium degradation level increases, while our _δ_^*t*^DTI test for differences in biological degradation maintains a very low false positive rate level of about 1% regardless of the amount of induced degradation. (**Figure 6a, 6b**).

**Figure 6.**
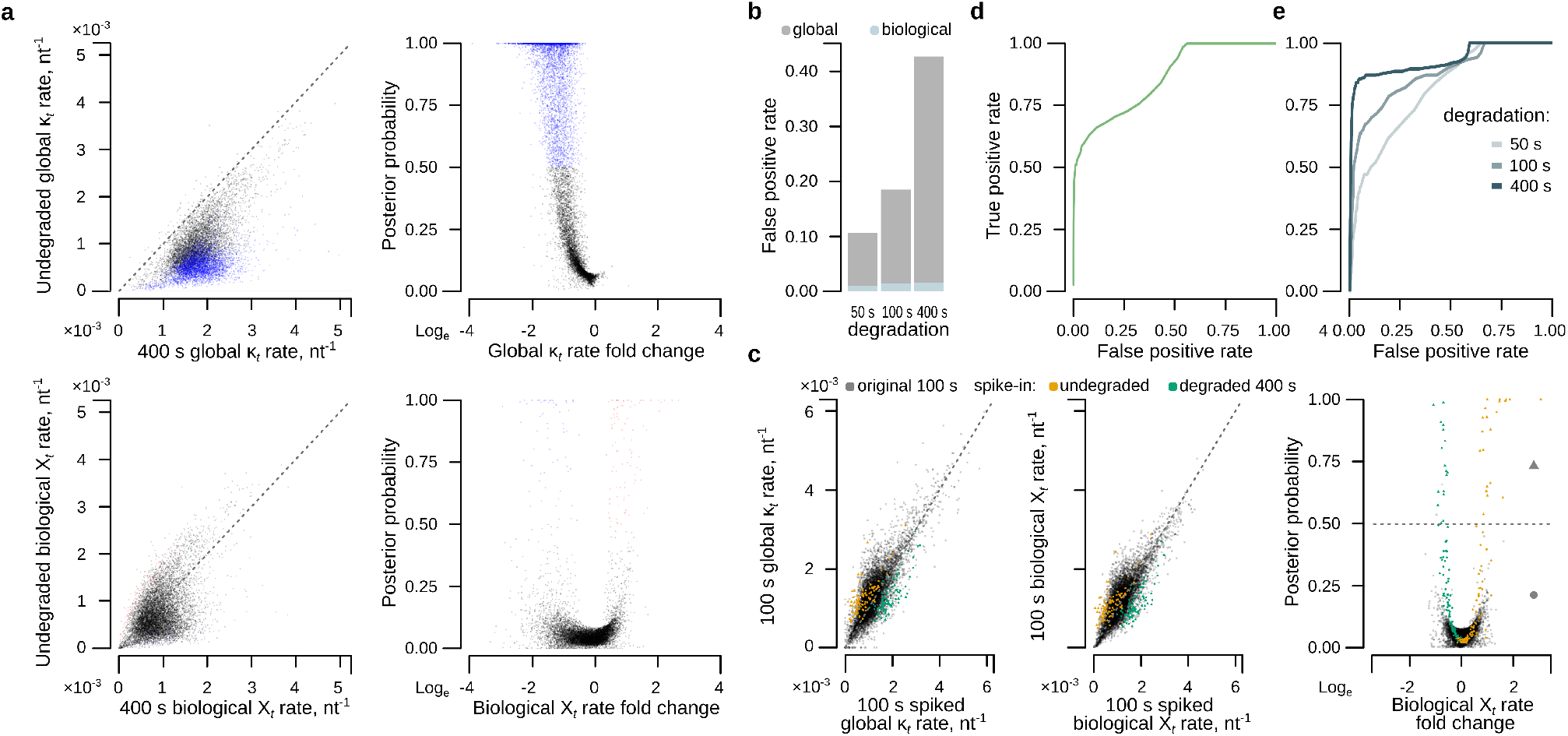
INDEGRA accurately deconvolves technical and biological components of the RNA degradation rate, and achieves high specificity and sensitivity in identifying differentially biologically degraded transcripts. **(a)** Comparison of the global fragmentation rates and distribution of significantly differentially degraded hits in the unadjusted data (top left panel) and same data with the technical component removed by INDEGRA (bottom left panel). Compared samples are HEK293 total RNA undegraded *versus* same RNA but subjected to magnesium ion-induced degradation at 75°C for 400 seconds. Scatter plots of the degradation difference posterior probability for the global (top) and technical component-removed (bottom) rates of the same data as in the left column (right column). **(b)** Comparison of significant false positive degradation hits identified between the undegraded control samples of HEK293 total RNA and samples with the same RNA subjected to magnesium ion-induced degradation of different intensity. Unadjusted hits from raw degradation rate measurements are shown in light blue, whereas residual technical component-removed hits are shown in dark blue. As a result of the correction, false discovery rate (FDR) is maintained at or below 1% regardless of the degradation intensity. **(c)** Validation of INDEGRA capacity to rectify biological degradation component away from the technical degradation using spike-in simulation with known ground truth. To achieve this, in one replicate of the 100-second magnesium ion-induced degradation samples, reads of 100 randomly selected transcripts were replaced with the reads of the same transcripts borrowed from 400-second degradation (green dots), whereas reads of 100 other randomly selected transcripts were replaced with the reads of the same transcripts borrowed from the undegraded sample (golden dots). Dot plots represent total (left) and biological (middle) fragmentation rates, and posterior probability of differential biological degradation per transcript with respect to biological degradation log-fold change(right). **(d)** ROC curve for data in (c). **(e)** ROC curve of alternative degradation spike-in simulation comparing technical replicate 1 of the undegraded condition with 100 transcripts having reads replaced by reads from the replicate 2 of a degraded condition, and replicate 1 of a degraded condition with 100 transcripts having reads replaced by reads from replicate 2 of the undegraded condition, for the 50-, 100- and 400-second degradation duration conditions

To validate the sensitivity of the test, we created two cross-spiked-in datasets. In the first, we compared the two replicates of a mild degraded condition (100 s) where in one replicate we replaced reads of 100 randomly selected transcripts by with reads of the same transcripts borrowed from the harshest degradation condition (400 s) and reads from another set of 100 randomly selected transcripts by with reads of the same transcripts borrowed from the undegraded condition. We show that at posterior probability threshold of 0.5, 95% of the identified hits are true positive (**Figure 6c**) and we obtain excellent receiver operating characteristic (ROC) curve (**Figure 6d**). In the second dataset, we compare one replicate of the undegraded condition to one replicate of a degraded condition (50 s, 100 s, 400 s) where reads from 100 random transcript of the undegraded replicate have been replaced with reads from the same transcripts from the other replicate of the same degradation condition, and reads from another 100 random transcrips of the degraded condition were replaced by reads from the same transcripts from the other replicate of the undegraded condition. In this scenario, as the degradation condition increases, comparing biological degradation between the samples becomes more challenging, while the spiked biological degradation difference of the replaced transcripts becomes more apparent. Here again, we achieve excellent ROC curves (**Figure 6e**).

### Correction of degradation bias in differential transcript abundance analysis

Finally, we discovered that DTI correlates highly with false positive differential transcript abundance (DTA; also translates to differential gene expression, DGE) hits (*R*^*2=*^0.846 for mouse and rat samples, while RIN correlates less, *R*^*2*^=0.721). This makes DTI an ideal dissection metric to identify and correct for those transcript counts that have been affected by the integrity differences. We thus implemented a LoWeSS regression-based correction to the non-linearity between the DTI-to-count function transcriptome-wide, as was previously done to correct for GC content bias^24^ (see **Materials and Methods, Figure 7a**). By applying this correction ahead of using standard DTA pipelines, INDEGRA balances samples depending on the technical degradation effects and provides a cleaned-up DTA (DGE) landscape (**Figure 7b**). For example, 310 DESeq2^25^ DTA hits were removed in samples with differences of 1.8 DTI (0.6 RIN) (**Figure 7b**), resulting in the corrected gene ontology (**Figure 7c**).

**Figure 7.**
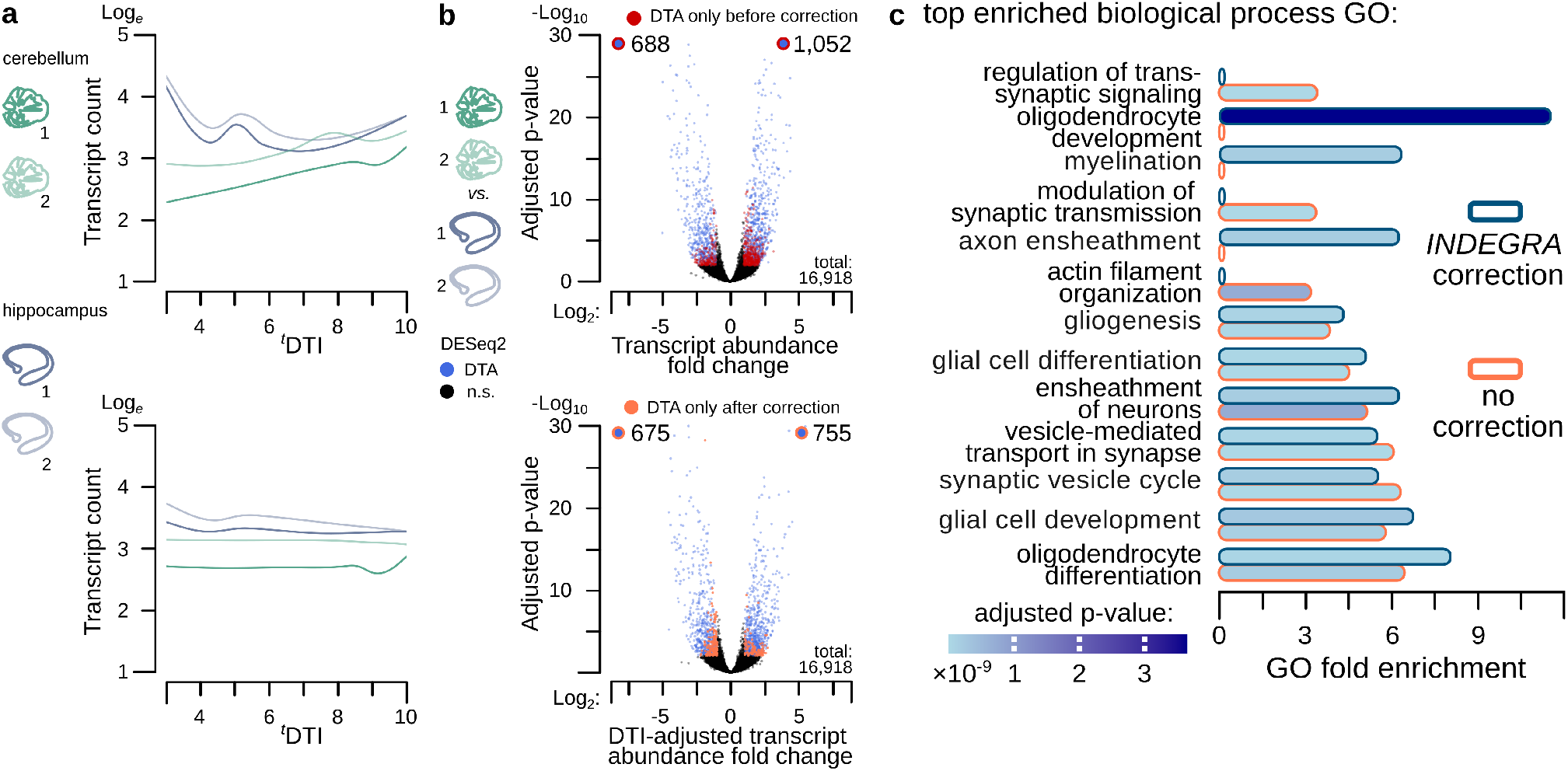
Application of INDEGRA to remove false differential gene expression hits. **(a)** Discovery of the ^*t*^DTI-dependent transcript count bias (top plot) and its correction using INDEGRA’s LoWeSS algorithm (bottom plot) shown with examples of biologically-replicated human total cerebellum and hippocampus RNA with different overall sample integrity. **(b)** Removing degradation-induced false differential transcript abundance (DTA) hits of DESeq2 with INDEGRA (same material as in (a)). **(c)** INDEGRA corrects gene ontology enrichment discovery for samples with different overall RNA degradation levels. Same material as in **(a**,**b)**

To validate this approach, we used the cross-spiked-in controlled HEK293 RNA degradation experiments to compare transcript abundance estimation between undegraded and several degraded conditions. We show that our approach significantly reduces the number of false positive hits, removing 25% to 60% of them even in the most degraded conditions (**Figures 8a, 8b**). To validate sensitivity of our approach, we created versions of the datasets with increased reads for a range of randomly selected transcripts representative of different length, abundance, ^*t*^DTI and isoformal complexity. First, comparing the undegraded to the 100 s degraded conditions, we successively randomly selected 100 transcripts with average relative abundance over all conditions within the 0.75 to 0.9 quantile range, 0.25 to 0.75 quantile range, and 0.1 to 0.25 quantile range. For those transcripts, we artificially increased their read counts by a factor 2.5 to 3.5 in either condition, and applied INDEGRA. In each case, INDEGRA significantly reduced the number of false positives while maintaining or improving the number of true positive hits compared to the standard uncorrected DTA approach (**Figures 8c, 8d**). We were able to achieve high sensitivity and specificity towards detecting true and suppressing false positives (**Figures 8e**). Second, we repeated this experiment comparing the undegraded to the different levels of degradation conditions, considering only the medium quantile range (0.25 to 0.75). Here again, we show that even in the most degraded conditions, INDEGRA achieves high specificity and sensitivity (**Figures 8e**).

**Figure 8.**
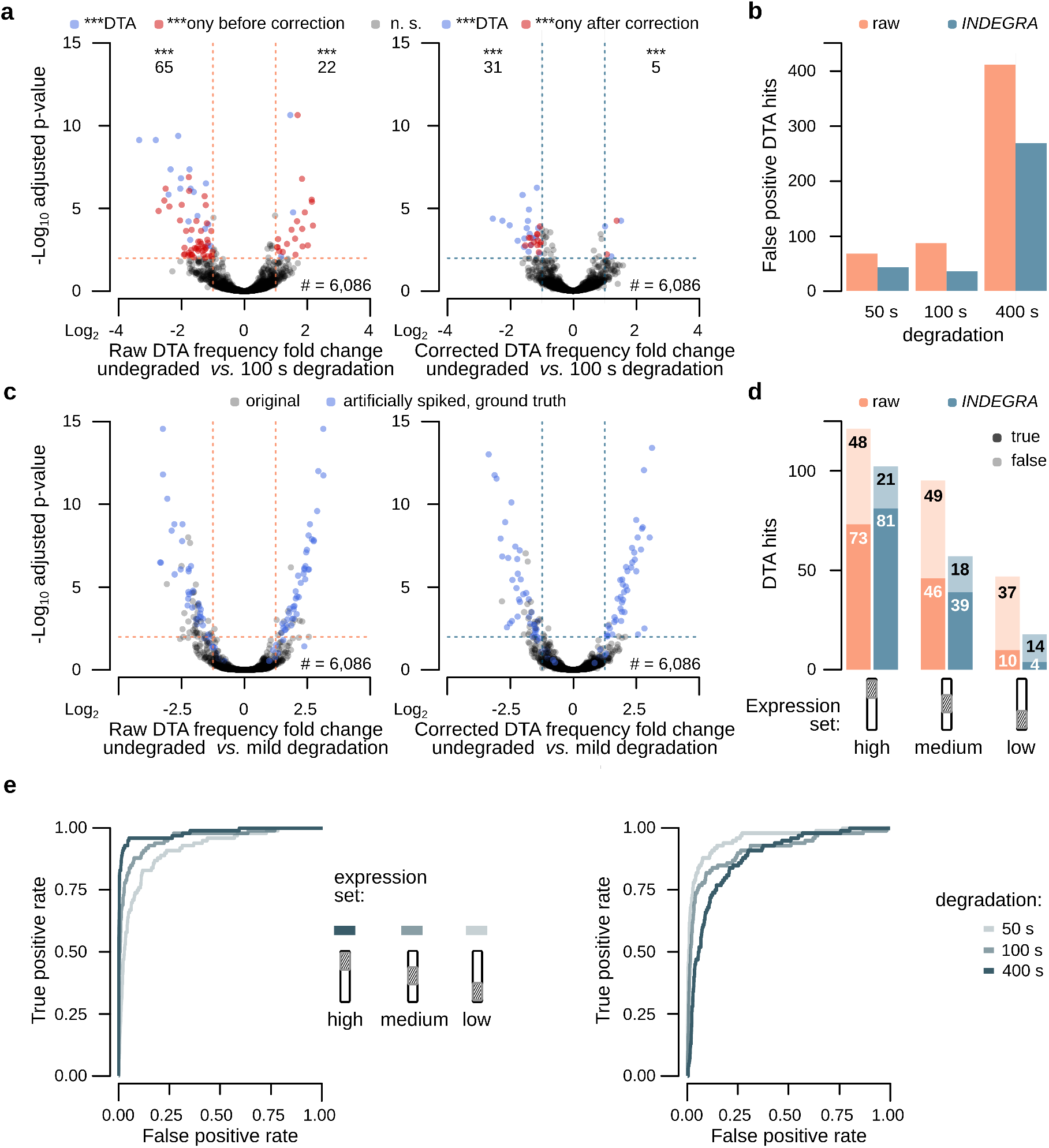
INDEGRA substantially and significantly reduces the number of false positive differential transcript abundance (DTA; also differential gene expression, DGE) hits while maintaining high specificity and sensitivity of the DTA detection. **(a)** Cleaning up DESeq2 DTA tests in controlled degradation experiments with INDEGRA. Shown are volcano plots comparing RNA from the undegraded HEK293 replicates (sample DTI of 7.00 and 6.77) to the same RNA upon 100-second magnesium ion-induced degradation replicates (sample DTI of 5.46 and 5.70). Without degradation correction, 87 false positive hits contaminated the outcome (top panel), whereas after correction only 36 false positive hits were found (bottom panel). **(b)** Reliable performance of INDEGRA in correcting DTA/DGE across a range of sample DTI differences. Number of false positive differential transcript abundance hits found with DESeq2 when comparing the undegraded HEK293 RNA samples to the same RNA subjected to the magnesium ion-induced degradation of increasing intensity. **(c)** Validation of false positive hit elimination in DGE/DTA by INDEGRA using differential transcript abundance simulation with known ground truth. 100 randomly selected transcripts with average relative abundance over all conditions in the 0.75 to 0.9 quantile range had read counts artificially increased by a factor of 2.5 to 3.5 in either condition. Shown are resultant volcano plots of DESeq2 output without degradation correction (left panel) and after degradation bias correction performed with INDEGRA (right panel) for comparison of undegraded HEK293 RNA samples to the same RNA subjected to the magnesium ion-induced degradation of 100 seconds. The ground truth hits are highlighted in light blue. **(d)** Reliable performance of INDEGRA in eliminating degradation difference-induced false positive DTA/DGE hits across different bands of relative transcript abundance. Number of true differentially expressed (DE, dark orange) and false positive (FP, light orange) transcripts identified by DESeq2 with and without INDEGRA correction in similar setting as in (c), but with the 100 transcripts chosen within the 0.75 to 0.9 quantile range (left), 0.25 to 0.75 quantile range (middle) and 0.1 to 0.25 quantile range (right), respectively. INDEGRA significantly and substantially suppresses the number of false positive hits and increases the number of true hits, reaching accuracy of 80% in the most challenging scenario (compared to 30% in the uncorrected analysis). **(e)** ROC curves for the outcome of experiments in (d) (left panel), and for the medium expressed set but comparing different levels of degradation (right panel), created by increasing p-value detection threshold from 0 to 1.

## Discussion

We present a robust, easy and accurate RNA degradation measure, the DTI, which can be used to characterize overall sample state as well as to reveal per-transcript integrity and degradation rate, accessibly based on DRS data. Calculated with INDEGRA, DTI highlights inter- and intra-transcript degradation variability, isolates RNA degradation from mapping uncertainties and links apparent degradation profile with the underlying absolute fragmentation rate of RNA. Using INDEGRA, it is possible to remove the majority of false differential transcript abundance hits resulting from the differences of overall sample integrity, while preserving transcript-specific degradation and stability differences. INDEGRA thus enables broad investigation of RNA turnover changes. INDEGRA directly plugs into the popular DTA/DGE instruments such as DESeq2, Limma-Voom^26^ and edgeR^27^. Our work facilitates unbiased RNA quantification in high-throughput data and streamlines comparisons of transcriptomes derived from different sources, across tissues, species and protocols.

## Materials and Methods

### RNA material and cell lines

HEK293 was originally purchased from the American Type Culture Collection (ATCC) and confirmed *via* short tandem repeat (STR) profiling with CellBank Australia^28^. The cells were grown in Dulbecco’s Modified Eagle Medium (DMEM) medium (Gibco) supplemented with 10% fetal bovine serum (FBS) and 1× antibiotic-antimycotic solution (Merck). Cells were cultured following the protocol previously described^28^. Briefly, HEK293 cell cultures were propagated in 1:3 splits, with replenishment of media every 4 days. Prior to cell collection, cells were propagated to a cell number of 1-2 million at cell viability and confluency of 90%. HEK293 cells were pelleted by gently spinning down the DMEM media containing the cells at 500 g for 5 minutes at room temperature (∼23°C) followed by two washes with ice cold phosphate buffered saline PBS (Thermo Fisher Scientific). The resultant cell pellets were stored at –80°C for all downstream applications. Total RNA extraction, random fragmentation-based degradation experiments (with spike-ins) and subsequent nanopore direct RNA sequencing were conducted using this HEK293 material.

### Purification of total RNA from HEK293

To isolate the total RNA fraction from HEK293 cells, we used PureLink RNA Mini Kit (Thermo Fisher Scientific), generally following the manufacturer’s protocol. 10 million cells of HEK293 the cell pellet obtained as described earlier were lysed in 500 μl of the denaturing lysis buffer as per manufacturer’s instruction. Cells were initially resuspended in the lysis buffer through pipetting and RNasin Plus (Promega) was immediately added to the cell lysate to a final concentration of 1 U/μl. Cell lysis was thoroughly facilitated by passing the cell lysate through a 27.5-gauge needle 6-8 times. Upon resting the cell suspension on ice for 5 minutes, the cell lysis was completed by passing the cell lysate through a 31-gauge needle for a total number of 6 cycles.

Following the cell lysis, the whole cell lysate was additionally resuspended by pipetting and then mixed with an equal volume of 80% v/v ethanol. The resultant mixture was transferred on top of a new silica binding column from the kit, assembled with its collection tube. The assembly was centrifuged for 30 seconds at 12,000 g. The protocol onwards exactly followed manufacturer’s instructions, and included the on-column DNase treatment. Briefly, 1 unit of TURBO DNase (Thermo Fisher Scientific) was added to the column matrix in ∼100 μl of the 1× TURBO DNase buffer (Thermo Fisher Scientific), and the column was then incubated for 15 minutes at room temperature (23°C), before proceeding to the wash steps. A series of wash steps with the Wash Buffers 1 and 2 was performed as recommended, and the purified RNA was eluted in ∼100 μl of deionised water. The initial quality and quantity of isolated RNA were assessed using Qubit Broad Range (BR) assay with Qubit spectrophotometer (Thermo Fisher Scientific). Where necessary, RNA quality, fragment size distribution and RIN metric were assessed by a Bioanalyzer 2100 (Agilent) run using RNA 6000 Nano chips. RNA was kept at –80°C until use.

### Synthesis of the reference RNAs

Reference RNAs i1, i3 and i4 were obtained by run-off *in vitro* transcription from plasmids encoding bacteriophage T7 RNA polymerase promoter followed by a unique artificial sequence covering the entire length of the desired RNAs (plasmids were ordered in sufficient amounts from GeneScript Biotech, are identical to those used by us earlier^29,30^ and contain synthetic RNA sequences with random 5-mers^31^. pUC57-mini-i-1, pUC57-mini-i-3 and pUC57-mini-i-4 DNA templates providing a spread over the RNA product lengths were generated by plasmid linearization with different restriction endonucleases, with pUC57-mini-i-1 set to yield 628 nt i1 RNA, pUC57-mini-i-3 – 1,072 nt i3 RNA and pUC57-mini-i-4 – 1,527 nt i4 RNA upon transcription.

#### Restriction enzyme digestion

Enzymes and buffers were purchased from New England Biolabs. pUC57-mini-i-1 was digested with AgeI, pUC57-mini-i-3 was digested with PstI, and pUC57-mini-i-4 was digested with HindIII. For each digestion reaction, 5 μg of each plasmid was mixed with 2 units of the corresponding restriction enzyme and supplemented with the enzyme buffer recommended by the supplier. The digestion reaction mixture was made to a total volume of 50 μl with deionized water, and was incubated at 37°C for 1 hour. To confirm the digestion outcome, samples were subjected to 1.5% w/v native agarose gel electrophoresis in 1× Tris-HEPES-EDTA buffer. Upon digestion confirmation, the digestion mixtures were made up to a total volume of 200 μl with deionized water and purified using 2× AMPure XP beads according to the supplier recommendation, upon which the purified DNA templates were eluted in 20 μl of deionized water and stored at –20°C until use.

#### Transcription reaction

Following the restriction digestion, template transcription was performed using HiScribe T7 High Yield RNA Kit (New England Biolabs) generally following manufacturer’s recommendations. For each transcription reaction, 2 μg of each template were mixed with nucleotide (NTP) Buffer Mix to a final concentration of 10 mM of each nucleotide, 5 mM DTT, 1 unit of RNasin Plus (Promega) and 2 units of T7 RNA polymerase in a total reaction volume of 40 μl. The transcription reaction mixture was then incubated at 37°C for 2 hours, after which the sample was diluted to 100 μl using deionized water supplied with 1× DNase I buffer (Thermo Fisher Scientific). 2 μl of 1 U/μl DNase I (Thermo Fisher Scientific) were added to the mixture, followed by incubation for 15 minutes at 37°C. To isolate the RNA, 2× volumes of AMPure XP SPRI bead suspension were added to this mixture, and binding was done for 5 minutes at 23°CC with periodic mixing. The binding was followed by two washing steps each with 1 ml of 80% v/v ethanol, as recommended by the bead supplier. The RNA was eluted with 30 μl of deionized water. The resultant purified spike-in i-type RNA samples were quality-controlled by a Bioanalyzer 2100 (Agilent) run using RNA 6000 Nano chips and their concentration additionally assessed with Qubit Broad Range (BR) assay using Qubit spectrophotometer (Thermo Fisher Scientific). RNA was kept at –80°C until use.

### Controlled magnesium ion-induced fragmentation of the i-type synthetic RNAs

i1, i3 and i4 RNAs were combined to produce an ‘equi-weight’ i-type master stock containing 20 μg of each RNA. This i-type stock mixture was then used to generate progressively more fragmented samples, with the fragmentation levels controlled by the time of exposure to high temperatures, in two independent replicates for each fragmentation level. In each experiment, i-type mixture containing total 5 μg of RNA was placed in a solution containing 50 mM HEPES and 5 mM MgCl_2_ based on deionized water. To ensure strict timing, concentration and temperature control, a heated-lid PCR machine and thinwall 200 μl PCR tubes were used. The PCR machine was set to 75°C for the exposure times of 50, 100, 200 and 400 seconds, and to 4°C immediately after that. After the fragmentation process, the samples were removed from the PCR machine and kept in ice for 2 minutes, after which the reaction mixture was supplemented with EDTA added to a final concentration of 5 mM. Two replicates of the control undegraded condition where no fragmentation was performed (no heating time) were also included in this setup. All samples were next diluted to 200 μl with deionized water. To remove any interfering reagent components or very short fragments for downstream applications from the fragmentation reaction mixture, a 2× AMPure XP bead purification was performed as described earlier. The RNA was eluted using deionized water in a final volume of 20 μl and assessed by a Bioanalyzer 2100 (Agilent) run using RNA 6000 Nano chips. RNA was kept at –80°C until use.

#### Bioanalyzer electrophoretic RNA mobility measurements

For all Bioanalyzer 2100 (Agilent) measurements of either quality check or physical degradation level assessment, standard RNA 6000 Nano kits and reagents, along with 2100 Expert software, were used. The Bioanalyzer image traces were used for qualitative validation of RNA mobility (and size as a function of that) after digestion and transcription reactions, and for calculating RIN values for RNA samples from HEK293 cell lines and human, mouse, rat, chicken, cow and dog tissues. High-quality image traces from Bioanalyzer were used for quantification of area-under-the-curve measurements using ImageJ software (https://imagej.net/) in degradation quantification after conducting the random fragmentation experiments on the reference i1, i3, i4 RNAs and HEK293 total RNA.

### *In vitro* polyadenylation of RNA

In order to mimic regular 3′-polyadenylated cytosolic mRNA and enable sequencing with the standard DRS protocol dependent on 3′ poly(A) priming, *in vitro* polyadenylation of i1, i3 and i4 RNAs was carried out. About 9 μg of the total RNA in 94 μl of deionized water or 25 mM HEPES-KOH (pH 7.6 at 25°C), 0.1 mM EDTA (HE) buffer were first denatured by incubating at 65°C for 3 minutes and immediately chilling in ice. The solution was then supplemented with 12 μl of *Escherichia coli* Poly(A) Polymerase buffer (New England Biolabs), 8 μl of 1 mM ATP and mixed. To the resultant solution, 3 μl of 40 U/μl RNasin Plus (Promega) and 3 μl of 5 U/μl *Escherichia coli* Poly(A) Polymerase (New England Biolabs) were added and mixed, and the resultant mixture was incubated at 37°C for 30 minutes. The reaction mixture was then brought to 200-300 μl with deionized water, and RNA was purified with 2× AMPure XP bead suspension as described earlier. Two independent replicates of each degradation condition and the control undegraded i-type RNA mixture samples were separately subjected to the *in vitro* polyadenylation reaction.

### HEK293 and i-type spike-in RNA random fragmentation

For the HEK293/i-type RNA spike-in fragmentation experiments, i1, i3 and i4 RNAs were individually polyadenylated as described earlier. Following this, the i1 and i3 poly(A) ^+^ RNAs have served as spike-ins for the random fragmentation experiment with HEK293 total RNA and were added from a premix before the fragmentation to independently control the fragmentation extent, whereas i4 RNA was added after the fragmentation process before conducting DRS library construction, to assess library, pore traversal and any other sequencing biases.

To produce the HEK293/i-type RNA spiked-in samples, 5 μg of the total RNA from HEK293 were mixed with 2.5 ng of each i1 and i3. If necessary, the mixture was proportionally scaled to produce sufficient material.

The controlled magnesium ion-induced fragmentation of the HEK293/i-type RNA spiked-in samples was conducted as described earlier, with all samples purified using 2× AMPure XP bead suspension. Finally, prior to conducting DRS library construction, 2.5 ng of i4 RNA was added to each sample.

### Direct RNA sequencing library preparation and loading

Flow cell priming, and library sequencing protocol were performed generally as previously described^28–30^. Nanopore sequencing was performed using an Oxford Nanopore Technologies’ MinION Mk1B with R9.4.1 flow cell for 72 hours. The default settings for the MinKNOW version 20.10.3 software were used and the SQK-RNA002 kit was selected. The flow cell priming, and library sequencing protocol were performed as follows. 5,000 ng of SPRI-purified RNA from each sample was used for each 2× library preparation within every replicate (all ONT recommended volumes doubled) with a direct RNA sequencing kit (SQK-RNA002) as supplied by ONT. SuperScript IV RNA Polymerase (Thermo Fisher Scientific) was used, RNA Control Standard (RCS) was omitted, and RNasin Plus (Promega) was included at 1 U/μl in all reaction solutions until the SPRI purification step after the reverse transcription reaction. The final adaptor-ligated sample was eluted in 40 μl.

### Basecalling and alignment

Reads from the i-type RNAs and HEK293 cell line were basecalled using Guppy version 6.4.6 and aligned to the corresponding reference transcriptome using minimap2^32^ with options ‘-ax map-ont - k 14’.

HEK293/i-type RNA spiked-in samples were aligned to a custom reference combining human transcriptome and i-type FASTA. Reads were then filtered to select the best alignment using samtools -F 2324^33^.

For the human, mouse, rat, chicken, cow and dog tissues, nanopore data were processed using Guppy version 5.0.7+2332e8d to basecall the nucleotide sequences, with parameters --flowcell FLO-MIN106 --kit SQK-RNA002 --device cuda:all:100% --compress_fastq --recursive -- num_callers 4 --gpu_runners_per_device 2 --chunks_per_runner 512 --chunk_size 1000.

To extract the transcriptome fasta sequence for each species, the program gffread (https://github.com/gpertea/gffread) was used. In the case of human data, GTF file (release version 102) was filtered to contain protein-coding transcripts, and transcriptome file was built using gffread.

Nanopore read sequences were mapped to the transcriptome of each species using minimap2 version 2.24-r11225 with parameters -ax map-ont -N 10.

### Preprocessing of the aligned reads

INDEGRA takes the transcriptome-aligned reads in the form of a BAM or SAM file produced with tools such as minimap2, and optionally the transcriptome reference GTF and alignment summary files, as the input.

By default, the INDEGRA code introduces an extra layer of filtering by discarding all reads with a maximum insertion length >80 nt, maximum deletion length >160 nt and cumulative softclip length on each end >200 nt (**Figure 2b**). Those thresholds are user-tunable.

For each isoform, 3′end mapping coordinate of the remaining reads are scanned using a cumulative sum sliding window of size 10 nt, to identify the most likely isoform 3′end position, which is termed ‘saturation point’ here. Reads with a 3′end position not within a 50 nt of the saturation point are then discarded (**Figure 2b**). Similarly, isoform 5′end positions are adjusted using the reads remaining after previous filtering steps reads as the most frequent read 5 ′ end position if it represents at least 10% of the reads start position, or left unadjusted based on the reference information otherwise (**Figure 2b**).

Retained reads are defined as ‘full length’ if their 5′end position is within a 15 nt window of the newly-defined isoform 5′end. Transcript length is then defined as the saturation point coordinate minus 5′end coordinate, any potential deletions and insertions are not counted.

Note that for performance and convenience purposes, the INDEGRA software can process several samples together in order to refine the isoforms’ 5′ and 3′ ends by combining all reads mapping to each isoform. Further estimation of degradation rates (see next paragraph) is performed on each sample separately.

### Testing for differential biological degradation

INDEGRA testing for differential biological degradation is performed in a Bayesian paradigm. A generative read-lengths distribution model conditioned on a set of random variables (one per transcript) is constructed, describing whether the two conditions have identical biological degradation (τ_*tj*_) or not. The technical degradation (α_*j*_) is accepted to occur after biological degradation, and therefore to affect all transcripts. Then, this model allows to deconvolve the global fragmentation rate κ_*tj*_ as 1 – (1 – τ_*tj*_) × (1 – α_*j*_), hence emphasizing that the technical degradation effect may not be linear.

Testing is then performed in a per-transcript manner by computing the posterior probability of the null hypothesis that biological degradation is the same between the two samples in question. Multiple testing control is performed *via* the prior probability of such event happening.

### Correcting transcript abundance from the degradation bias

An approach similar to described previously for GC content correction is used^24^. The log read count of each transcript is regressed by their ^*t*^DTI value. The regressing function is then removed from the original counts while median is added to preserve the library scale. Such correction is then provided as an offset to the downstream differential expression tools such as DESeq2, edgeR or limma-voom, enabling a universal, non-destructive adjustment.

### The INDEGRA software

The INDEGRA suit is hosted at https://github.com/Arnaroo/INDEGRA and comprises three programs.

To compute the DTI, a python code extracts all necessary information from the BAM files and the optional supplementary information such as GTF or sequencing summary file, and outputs a per-transcript summary CSV file with ^*t*^DTI information and necessary statistics for any downstream analysis. Optionally, the python code may return a cleaned-up BAM files with all discarded reads removed.

Differential degradation and differential transcript abundance (gene expression) analyses are then performed *via* R codes. The differential degradation script can processes two samples at a time, and returns the posterior probability of same biological degradation as well as posterior expectation of biological degradation rate in each condition.

Differential abundance (gene expression) correction can then be accessed and can process together as many samples as desired. Corrected differential transcript abundance (gene expression) is provided as an offset matrix to feed into the downstream software. Examples and pre-made scripts for DeSeq2, edgeR and limma-voom are included, however any other software can be used as well. More details and instructions can be found at https://github.com/Arnaroo/INDEGRA GitHub.

### Application of INDEGRA across datasets

The INDEGRA software was applied using default parameters on the set of spiked-in HEK293 samples combined. Differential degradation was then evaluated using replicates of each condition with the most reads, while differential transcript abundance (gene expression) analysis was performed using all samples together.

For the 63 samples of different species and tissues, the INDEGRA software was applied using default parameters and processing of all samples from a species together. Posterior expectation of biological degradation rates was then obtained per-sample using the differential degradation script, and then combined. Heatmaps of degradation rates were plotted based on these data using pheatmap R package.

Differential transcript abundance correction was only performed on paired conditions simultaneously, and the offset matrix was provided to DeSeq2, edgeR and limma-voom run with otherwise standard parameters.

## Data availability

The data used in this manuscript are all downloaded from publicly available data sources, or made publicly available if generated by the authors. Specifically, PRJNA1196015 was generated for the purposes of this study and PRJNA1188790 was generated in a parallel work^34^. All relevant information about data is described in the Methods.

## Code availability

The INDEGRA package contains all code required to perform the global and per-transcript DTI calculation, separation of the biological and technical degradation as well as instructions to install and run the software, and is publicly available for free use at a maintained repository over https://github.com/Arnaroo/INDEGRA. INDEGRA version used at the time of publishing is also available as a static Figshare item https://doi.org/10.6084/m9.figshare.27689616. Core protocols used in the work are also available at protocols.io under https://dx.doi.org/10.17504/protocols.io.261gerdyol47/v1

## References

1. Mattick, J. S. RNA out of the mist. Trends in Genetics 39, 187–207 (2023).

2. Houseley, J. & Tollervey, D. The Many Pathways of RNA Degradation. Cell 136, 763–776 (2009).

3. Arraiano, C. M. et al. The critical role of RNA processing and degradation in the control of gene expression. FEMS Microbiology Reviews 34, 883–923 (2010).

4. Ottens, F., Efstathiou, S. & Hoppe, T. Cutting through the stress: RNA decay pathways at the endoplasmic reticulum. Trends in Cell Biology 0, (2023).

5. Tuck, A. C. et al. Mammalian RNA Decay Pathways Are Highly Specialized and Widely Linked to Translation. Mol Cell 77, 1222–1236.e13 (2020).

6. Chen, C. A., Ezzeddine, N. & Shyu, A. Chapter 17 Messenger RNA Half-Life Measurements in Mammalian Cells. in Methods in Enzymology vol. 448 335–357 (Academic Press, 2008).

7. Czarnocka-Cieciura, A. et al. mRNA decay can be uncoupled from deadenylation during stress response. 2023.01.20.524924 Preprint at 10.1101/2023.01.20.524924 (2023).

8. Tosar, J. P., Witwer, K. & Cayota, A. Revisiting extracellular RNA release, processing, and function. Trends Biochem Sci 46, 438–445 (2021).

9. Łabno, A., Tomecki, R. & Dziembowski, A. Cytoplasmic RNA decay pathways - Enzymes and mechanisms. Biochimica et Biophysica Acta (BBA) - Molecular Cell Research 1863, 3125–3147 (2016).

10. Brothers, W. R., Ali, F., Kajjo, S. & Fabian, M. R. The EDC4-XRN1 interaction controls P-body dynamics to link mRNA decapping with decay. EMBO J 42, e113933 (2023).

11. Kurosaki, T. & Maquat, L. E. Nonsense-mediated mRNA decay in humans at a glance. Journal of Cell Science 129, 461–467 (2016).

12. Karasik, A., Lorenzi, H. A., DePass, A. V. & Guydosh, N. R. Endonucleolytic RNA cleavage drives changes in gene expression during the innate immune response. Cell Reports 43, (2024).

13. Yamagami, R., Sieg, J. P. & Bevilacqua, P. C. Functional Roles of Chelated Magnesium Ions in RNA Folding and Function. Biochemistry 60, 2374 (2021).

14. Prawer, Y. D. J., Gleeson, J., De Paoli-Iseppi, R. & Clark, M. B. Pervasive effects of RNA degradation on Nanopore direct RNA sequencing. NAR Genomics and Bioinformatics 5, lqad060 (2023).

15. Schroeder, A. et al. The RIN: an RNA integrity number for assigning integrity values to RNA measurements. BMC Molecular Biology 7, 3 (2006).

16. Matsubara, T. et al. DV200 Index for Assessing RNA Integrity in Next-Generation Sequencing. BioMed Research International 2020, 9349132 (2020).

17. Wang, L. et al. Measure transcript integrity using RNA-seq data. BMC Bioinformatics 17, 58 (2016).

18. Swanson, G. M., Estill, M. S. & Krawetz, S. A. The transcript integrity index (TII) provides a standard measure of sperm RNA. Systems Biology in Reproductive Medicine 68, 258–271 (2022).

19. Feng, H., Zhang, X. & Zhang, C. mRIN for direct assessment of genome-wide and gene-specific mRNA integrity from large-scale RNA-sequencing data. Nat Commun 6, 7816 (2015).

20. Garalde, D. R. et al. Highly parallel direct RNA sequencing on an array of nanopores. Nat Methods 15, 201–206 (2018).

21. Depledge, D. P. et al. Direct RNA sequencing on nanopore arrays redefines the transcriptional complexity of a viral pathogen. Nat Commun 10, 754 (2019).

22. Lee, J. Generalized Bernoulli process with long-range dependence and fractional binomial distribution. Dependence Modeling 9, 1–12 (2021).

23. Benjamini, Y. & Hochberg, Y. On the Adaptive Control of the False Discovery Rate in Multiple Testing With Independent Statistics. Journal of Educational and Behavioral Statistics 25, 60–83 (2000).

24. Risso, D., Schwartz, K., Sherlock, G. & Dudoit, S. GC-content normalization for RNA-Seq data. BMC Bioinformatics 12, 480 (2011).

25. Love, M. I., Huber, W. & Anders, S. Moderated estimation of fold change and dispersion for RNA-seq data with DESeq2. Genome Biology 15, 550 (2014).

26. Ritchie, M. E. et al. limma powers differential expression analyses for RNA-sequencing and microarray studies. Nucleic Acids Research 43, e47–e47 (2015).

27. Robinson, M. D., McCarthy, D. J. & Smyth, G. K. edgeR : a Bioconductor package for differential expression analysis of digital gene expression data. Bioinformatics 26, 139–140 (2010).

28. Sneddon, A. et al. Biochemical-free enrichment or depletion of RNA classes in real-time during direct RNA sequencing with RISER. Nat Commun 15, 4422 (2024).

29. Horvath, A. et al. Comprehensive translational profiling and STE AI uncover rapid control of protein biosynthesis during cell stress. Nucleic Acids Research 52, 7925–7946 (2024).

30. Acera Mateos, P. et al. Prediction of m6A and m5C at single-molecule resolution reveals a transcriptome-wide co-occurrence of RNA modifications. Nat Commun 15, 3899 (2024).

31. Liu, H. et al. Accurate detection of m6A RNA modifications in native RNA sequences. Nat Commun 10, 4079 (2019).

32. Li, H. Minimap2: pairwise alignment for nucleotide sequences. Bioinformatics 34, 3094–3100 (2018).

33. Danecek, P. et al. Twelve years of SAMtools and BCFtools. GigaScience 10, giab008 (2021).

34. Santos-Rodriguez, G. et al. The conserved landscape of RNA modifications and transcript diversity across mammalian evolution. 2024.11.24.624934 bioRxiv Preprint at 10.1101/2024.11.24.624934 (2024).

